# Adaptation and phenotypic diversification of *Bacillus thuringiensis* 407 biofilm are accompanied by a fuzzy spreader morphotype

**DOI:** 10.1101/2021.09.03.458824

**Authors:** Yicen Lin, Gergely Maróti, Mikael Lenz Strube, Ákos T. Kovács

## Abstract

*Bacillus cereus* group (*Bacillus cereus sensu lato*) has a diverse ecology, including various species that produce biofilms on abiotic and biotic surfaces. While genetic and morphological diversification enable the adaptation of multicellular communities, this area remains largely unknown in the *Bacillus cereus* group. In this work, we dissected the experimental evolution of *Bacillus thuringiensis* 407 Cry-during continuous recolonization of plastic beads. We observed the evolution of a distinct colony morphotype that we named fuzzy spreader (FS) variant. Most multicellular traits of the FS variant displayed higher competitive ability versus the ancestral strain, suggesting an important role for diversification in the adaptation of *B. thuringiensis* to the biofilm lifestyle. Further genetic characterization of FS variant revealed the disruption of a guanylyltransferase gene by an insertion sequence (IS) element, which could be similarly observed in the genome of a natural isolate. The evolved FS and the deletion mutant in the guanylyltransferase gene (Bt407Δ*rfbM*) displayed similarly altered aggregation and hydrophobicity compared to the ancestor strain, suggesting that adaptation process highly depends on the physical adhesive forces.

## Introduction

Multicellularity, assemblage of differentiated cells, is one of the novel evolutionary strategies among microbes. Although a well-accepted concept for eukaryotes, it took several decades to generally acknowledge that nearly all bacteria are capable of multicellular behaviors^1^. Yet, bacteria are nowadays broadly harnessed to study evolution of multicellularity due to their simpler characteristics. Among general classes of multicellular bacteria, aggregation is regarded as one of the most critical class. Embedded within the extracellular matrix (ECM) of biofilms, aggregation mediated cell–cell adhesion provides fitness advantage for bacteria in various ways^2^. For instance, by holding cells together, ECM prevents the noxious influence of external toxic substances like antimicrobial compounds and enables community members to share enzymes and retain liquid repellency^3,4^. In *Myxococcus xanthus*, ECM is responsible for coordinated movement such as swarming motility by building cell collectives and triggering pilus retraction^5^. The relatively large size of multicellular aggregates might also confer a selective advantage compared with individual cells when facing predation^6^. More importantly, in environments where resources are inadequate for unicellular growth, aggregation can support direct access to nutrients produced by neighboring cells^7–9^. In analogy to nutrient transport in the veins of the animal body, secreted compounds can be dispersed through channels within structured biofilms. Likewise, multicellular aggregates also have predominant fitness advantage over solitary cells on plant leaves^10,11^. The advantages of multicellularity are plentiful, primarily due to the larger microbial biomass created through physical adhesion. These benefits can offset the negative effects that multicellularity brings, such as impaired motility and increased demand for food resources owing to higher cell density^2,12,13^.

Experimental evolution studies have been utilized to explore bacterial adaptive diversification for decades. One of the simplest model uses glass tubes culturing *Pseudomonas fluorescens* under static condition^14^. Driven by spatial heterogeneity and competition for vacant niches, *P. fluorescens* rapidly diversifies into three distinct colony morphotypes. The wrinkled-spreader morphotype with multicellular characteristics forms a self-supporting mat at the air-liquid surface. Owing to its simplicity and reproducibility, this static microcosm serves as a model to study evolutionary diversification that eventually expands our ecological and genetic understanding of *P.fluorescens*^15–18^. Additional simple setups have also been successfully applied to study evolution outcome, such as colonies on solid agar plates^19,20^, submerged biofilms in microtiter plates^21^, and *in silico* models simulating static systems^22,23^. Despite the variations among these experimental setups, the generally applied spatially structured environments provide ecological opportunities in the form of distinct niches, where the diversifying selection is driven by resource competition. When such spatial structure is destroyed by shaking, the diversification is also eliminated^14^. However, when a vacant niche is constructed, even in shaking condition, heterogeneity within biofilms can still provide ecological opportunities for adaptive diversification, as demonstrated in a bead model^24^. While the evolution of biofilms has been mostly studied in Gram-negative bacteria, including *Pseudomonas spp*. and *Burkholderia cenocepacia*, much less attention has been given to Gram-positive bacteria. Within Gram-positive and spore-forming bacteria, *Bacillus subtilis* has been exploited to reveal that adaptive specialization readily occur under aerated cultivation via mutations that influence the regulation of biofilm development^25^. Focusing on the air-liquid floating biofilm, called pellicle that create a highly structured environment, pellicles of *B. subtilis* underwent significant evolutionary diversification after ca. 200 generations, including an exploitative interactions among different evolved morphotypes^26^. Similarly, reduced spatial heterogeneity or hampered motility can also select for higher matrix production^27^. Recent works started to exploit *Bacilli* to understand how colonization of a plant host influence bacterial evolution^28–30^.

*Bacillus cereus* group species are widely observed in soil samples. A consensus view is that *B. cereus sensu lato* can proliferate either within animal hosts or in the rhizosphere, exhibiting either pathogenic or symbiotic lifestyles^31^. Switching to multicellularity is one of the key strategies to thrive in these diverse niches. For example, in the guts of arthropods, *B. cereus* can grow by creating multicellular structures^32,33^. Another impressive example nicely demonstrated how *B. cereus* grew in soil-extracted organic matter medium by employing a growth of multicellular mode embedded in extracellular matrix^34^. Collectively, these examples indicate a conserved capacity among *B. cereus* species to grow in multicellular structures like cell aggregates and chains.

Despite the considerable amount of literature on the multicellular growth of *B. cereus* species, there has been little evidence of experimental evolution and ecological benefits of multicellular structures in these bacteria. In this study, we seek to explore how *Bacillus thuringiensis* (*Bt*) adapt to cycles of biofilm formation and illustrate that *Bt* could adopt a multicellular lifestyle to retain fitness advantage in a bead colonization model. This model routinely selects for cells colonizing the surface of plastic beads and subsequently dispersing from it and therefore it creates a simplified selection for bacterial life cycle.

## Results

### Evolution of biofilm on plastic beads is accompanied by diversification of Bt4O7 into a distinct morphotype

To test the adaptive diversification of *B. thuringiensis* 407 Cry-(Bt407), six independent populations were experimentally evolved as biofilms. We utilized the renowned nylon bead-based biofilm experimental evolution system developed by Poltak and Cooper^24^, but included three beads in each new inoculation steps: one colonized bead from the previous step and two sterile beads that were subsequently used for the next transfer or to determine the bacterial cell counts (Fig. 1A). Around every three transfers, biofilm developed on one of the beads was dispersed using sonication, and cells were plated on lysogeny broth (LB) agar plates for colony forming unit (CFU) counting. Biofilm productivity (i.e. cell counts of the biofilms developed on the beads) increased gradually compared with the initial inoculum until transfer 31, after which a decrease was observed (Fig. S1). Using a regression analysis, robust linear correlation was found between the timeframe and biofilm productivity (*R*^2^ of 0.77, *p* < 0.01, Fig. S2) until transfer 31 (with removing outlier at transfer 25). At the end, all six biofilm populations displayed significantly enhanced biofilm production, suggesting the bead biofilm model serves as a great tool to select for biofilm-forming lineages (Fig. 1B). On the contrary, six experimentally evolved populations that were continuously cultivated under planktonic conditions did not exhibit significant difference in their biofilm formation ability compared with the ancestor (Fig. S3).

**Fig. 1.**
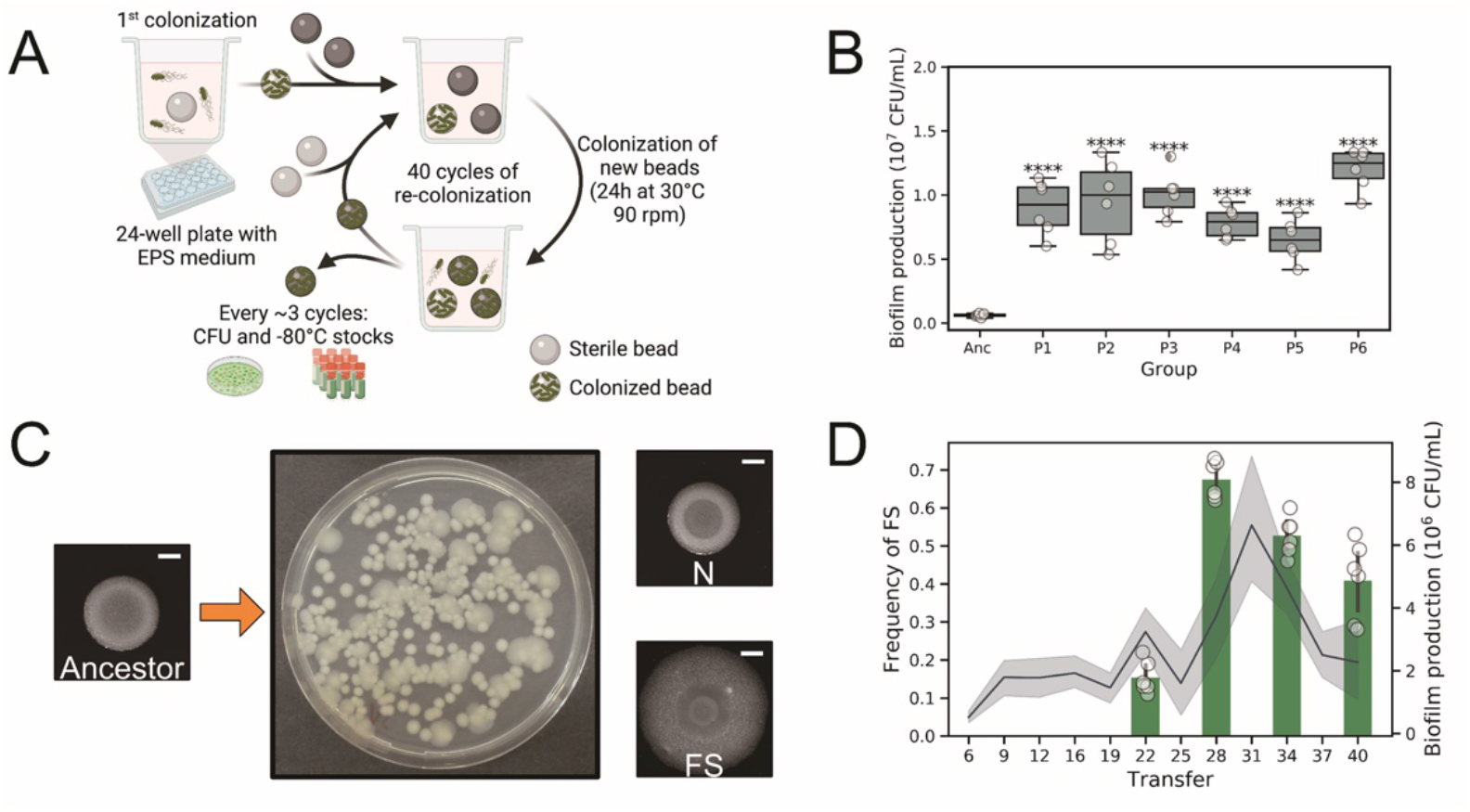
Evolution of nylon bead associated biofilms of Bt407 diversified into a distinct morphotype. (A) Schematic representation of the bead-based experimental evolution setup. (B) Enhanced biofilm productions revealed by CFU analysis of all final evolved populations compared with the ancestor (n = 6 biologically independent samples). Boxes indicate Q1–Q3, lines indicate the median, and bars span from max to min. Asterisks indicate significant differences between each group and the ancestor (\*\*\*\**p* <0.0001; One-way ANOVA followed by Dunnett’s multiple comparison tests). (C) Distinct colony morphotypes of the evolved variants were evidenced on LB agar medium. Scale bars indicate 5 mm. (D) Bar plot represents the relative frequencies of the FS morphotype (n=6). Line plot represents the average biofilm productivity along the experimental evolution. Error bars and shaded area indicate the standard error of the mean in the bar chart and line chart, respectively.

In structured environments, microbes can diversify into phenotypically different variants, which are identified as morphotypes as they exhibit distinguishable colony morphologies. Biofilms on the beads are regarded as a classical example of such a structured environment. In all evolved biofilm populations, a distinct morphotype, demonstrating large and surface colonizing colony with relatively translucent edges (fuzzy spreader), was identified on LB agar plates (Fig. 1C). On the contrary, no such evolved variants were observed in planktonic evolved populations. The other colony morphotype was indistinguishable from the ancestor and therefore was termed as normal (N). To unravel the evolutionary history of phenotypic diversification and determine the frequencies of FS in all populations, fractions of frozen stocks from selected evolutionary time points were serially diluted in 0.9% sodium chloride solutions and plated onto LB agar. FS morphotypes were first detected around transfer 22 with a maximum frequency at transfer 28. Notably, the increased frequencies of FS coincided with the maximum biofilm productivity, implying FS morphotype may act as a biofilm-specialist in the bacterial populations (Fig. 1D).

### Phenotypic variation affects preference of biofilm formation in Bt407 evolved isolates

To further elucidate the phenotypic variations among morphotypes, we compared fundamental cellular differentiation properties of the derived two morphotypes from population 1 (P1) testing biofilm formation ability and motility (Fig. 2 and Fig. S4). The FS morphotype displayed increased Congo red uptake, suggesting that the FS variant had enhanced exopolysaccharide production compared with the ancestor. In addition to a slightly increased wrinkleality, the FS colonies demonstrated enhanced spreading motility on both EPS (29mm ± 4mm) and Trb media plates (33mm ± 6mm) with 0.7% of agar. On EPS agar, normal variants (11mm ± 3mm) and the ancestor (15mm ± 4mm) demonstrated similar surface motility on Trb agar, no significant difference was found between normal variants (15mm ± 5mm) and the ancestor (14mm ± 4mm) either. Intriguingly, FS variants displayed dendritic (branched) spreading, especially on EPS medium. Dendritic formation of colonies depends on nutrient resource and is promoted under lower nutrient conditions. The dendritic colony of the FS variant might represent an alternative surface translocation strategy possibly contributing to fitness during the selection process. While the N morphotype showed comparable colony morphology to the ancestor on LB agar medium, it displayed increased uptake of Congo red and decreased colony spreading ability on both spreading plates, suggesting distinct influence on biofilm development compared to the ancestor. These properties of FS and N morphotypes from population 1 could also be demonstrated for isolates from the other five populations (Fig. S4), highlighting parallel evolution in all representative populations.

**Fig. 2.**
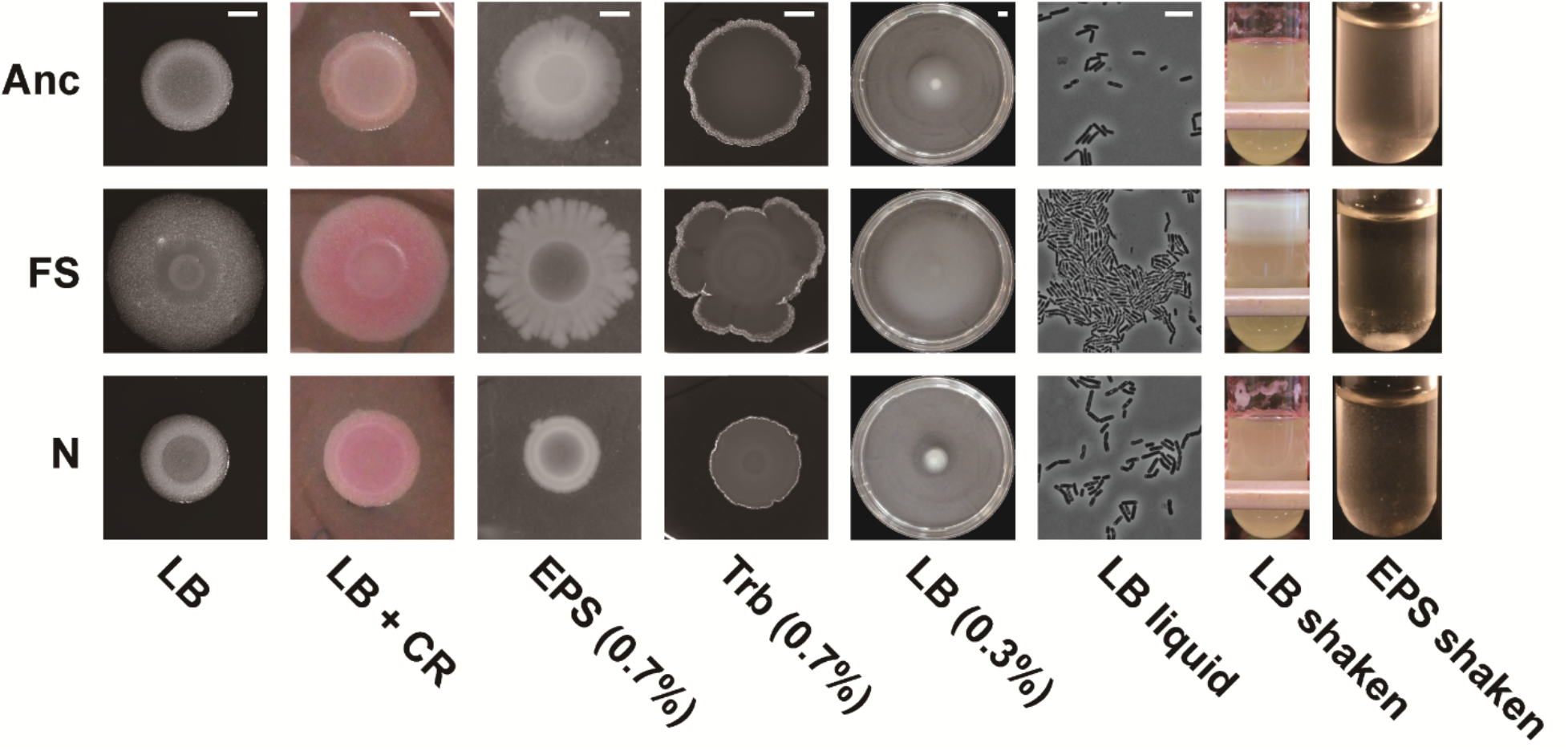
Phenotypic determination of the evolved variants. From left to right: colony morphologies, Congo red uptake, swarming motility on EPS and Trb medium, swimming motility, cellular morphologies in LB liquid, biofilm formation in LB and EPS medium in shaken cultures. Anc, FS and N indicate the ancestor, FS and N morph variants, respectively. Scale bars in colony and surface motility images indicate 5mm and that in single cellular morphologies indicates 10 μm across all groups.

Motivated by the general positive correlation between Congo red binding and biofilm formation^35,36^, biofilm formation of the morphotypes was tested in shaken LB cultures. Both evolved variants exhibited increased biofilm formation on the tube walls, especially the FS variant. In the nutrition limited EPS medium, the FS variant largely formed non-attached aggregates in the liquid fraction, further suggesting that the selection led to multicellularity by the FS morphotype. The N variant showed intermediate biofilm-related phenotypes. Besides, the cells of FS morphotype exhibited dense and aggregated behavior as observed under microscope, again suggesting an elevated biofilm formation ability. Taken together, the FS variant exhibited dramatically higher degree of multicellularity compared with the ancestor, while the N morphotypes might serve as a stable member of the microbial community contributing to a different function within the biofilm.

### Evolution of synergistic association of evolved isolates in bead biofilms

Bacterial evolution often involves trade-off between planktonic growth and biofilm formation. Importantly, the theory that trade-off in competition-colonization facilitates adaptive radiation is widely accepted. To reveal any trade-off in the evolved morphotypes, the variants were quantitatively tested in planktonic and biofilm states and compared to the ancestor strain. Both morphotypes exhibited reduced growth in planktonic cultures compared with the ancestor, suggesting that higher fitness in biofilm formation is at the expense of planktonic doubling time (Fig. 3A). Nonetheless, when the two morphotypes incubated together, the growth rate of the well-mixed liquid cultures was restored to the level of ancestor (Fig. 3A), which might point towards synergistic metabolic connection between the two evolved variants.

**Fig. 3.**
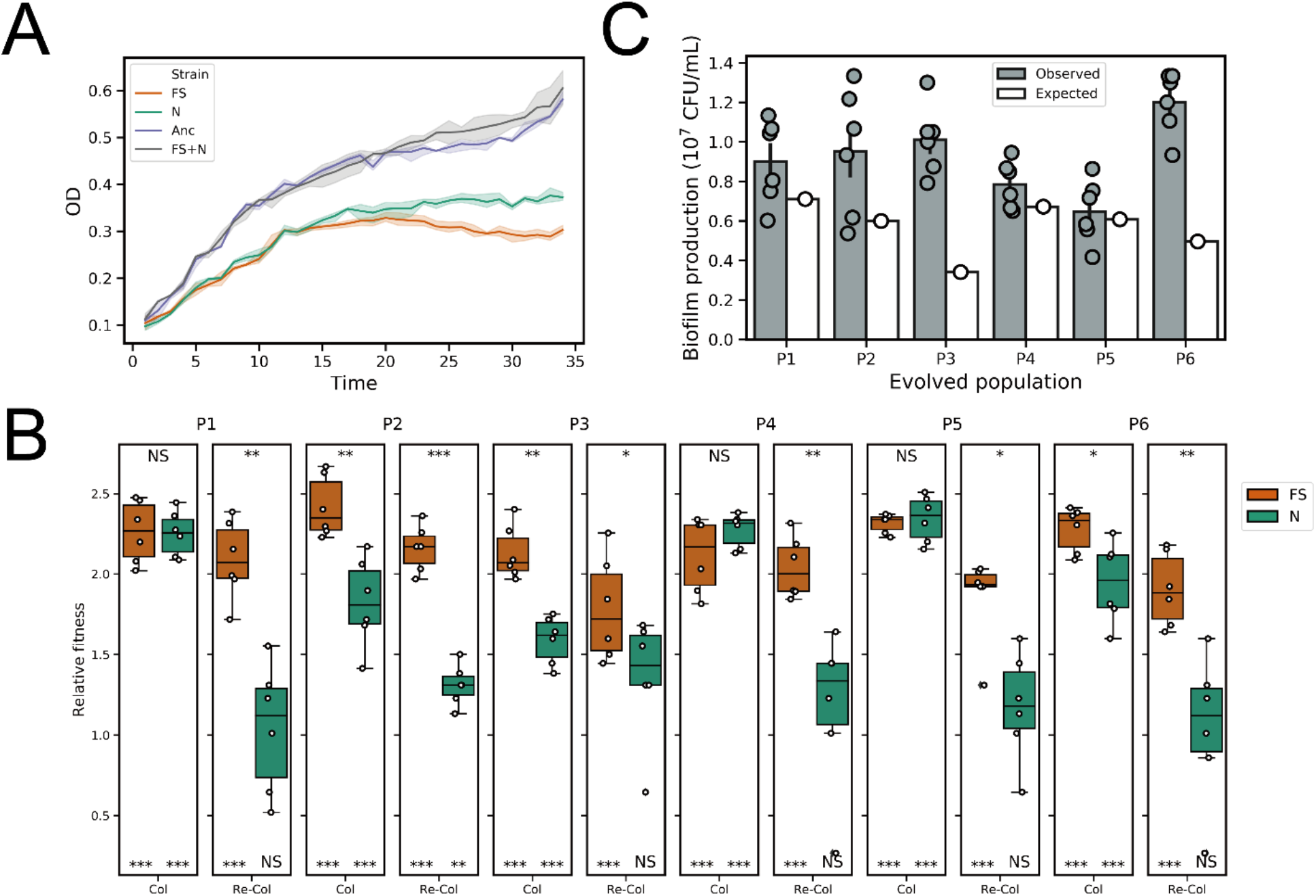
Morphotype fitness in planktonic and biofilm forming conditions. (A) Growth properties of the evolved variants and the ancestor (n = 6). (B) Relative fitness of the evolved variants in the bead biofilms compared with the ancestor. Columns indicate colonization and recolonization of evolved populations, respectively (P1-P6). Boxes indicate Q1–Q3, lines indicate the median, and bars span from max to min. Asterisks at the top indicate significant differences between FS and N morph variants. Asterisks at the bottom indicate significant differences between each group and the ancestor (\**p* <0.05, \*\**p* <0.01, \*\*\**p* <0.001; One-way ANOVA followed by Dunnett’s multiple comparisons tests). (C) Observed and expected biofilm productions of six evolved populations (P1-P6). Expected productivities were calculated as the product of the proportion of the evolved variant in each population and their productivities in monocultures. For observed biofilm productions, error bars indicate standard error of the mean of independent biological samples (n = 6).

Relative fitness of representative morphotypes from all evolved populations was calculated as the ratio of Malthusian parameter described by Lenski *et al*.^37^. Testing the evolved variants from all 6 populations confirmed increased fitness compared to the ancestor when cultures colonized the nylon beads. All FS variants exhibited stable and robust fitness advantage over the ancestor, relative fitness ranging from 2.11 to 2.41 in average (Fig. 3B), while the N variants also had increased fitness, but displaying higher variability, 1.58 to 2.34 in average. When two-cycle bead colonization was tested, termed re-colonization, the fitness advantage of FS variants remained high compared to the ancestor (1.78-2.15). On the contrary, the relative fitness of N morphotype during the bead re-colonization was comparable to the ancestor. The statistical comparison between FS and N variants further confirmed the latter ones had reduced re-colonization fitness. Notably, while the FS and N isolates from 3 populations (P2, P3, and P6) showed significantly different fitness during colonization, isolates from the other 3 populations exhibited comparable fitness, suggesting that the N variants across evolved populations may not have acquired similar fitness advantage and phenotypic adaptation, unlike FS variants. Noteworthy, re-colonizing ancestor cells produced dramatically less biofilms than colonizing cells (*p* < 0.01, Fig. S5), indicating a strong selection bottleneck for immigrating cells from old biofilms to new ones.

Finally, using the determined biofilm yield of monocultures, the expected productivities in the final mix cultures were calculated based on their frequencies in the mix as previously described^24^. Not surprisingly, the expected yield of the mix cultures was lower compared with the observed productivities (Fig. 3C), which was in accordance with previous literature^24,26,38^. Notably, the difference between the predicted and observed productivity was higher in populations P2, P3, and P6, in which the FS and N variants exhibited significantly different colonization fitness values.

### Multicellular characteristics of evolved FS morphotypes confer ecological benefits

The spatial organization of evolved variants within biofilms influences the relative fitness of the population and effects competitive or cooperative traits^24,39–41^. Therefore, to further explore the microbial populations, variants of population P1 were fluorescently labeled along with the ancestor, and the spatial distribution of each morphotype and ancestor were imaged. Confocal laser scanning microscopy (CLSM) imaging demonstrated that FS variants gained a substantial competitive advantage over normal isolates and the ancestor, exhibiting large multicellular clusters in the biofilms (Fig. 4). While the FS variant strongly outcompeted the ancestor, when co-cultured with the N variant, the FS morphotype became evenly distributed in the biofilm, producing smaller aggregates compared with the FS-ancestor co-cultures. Notably, the normal variants occupied the bottom layer of the biofilms, while the FS morphotype resided the upper region, suggesting a spatial arrangement among the evolved variants. The spatial diversification of the morphotypes might explain the increased total biofilm yield compared with the expected calculation. Furthermore, the quantification of CFU within co-cultured biofilms also confirmed the competitive advantage of FS variants.

**Fig. 4.**
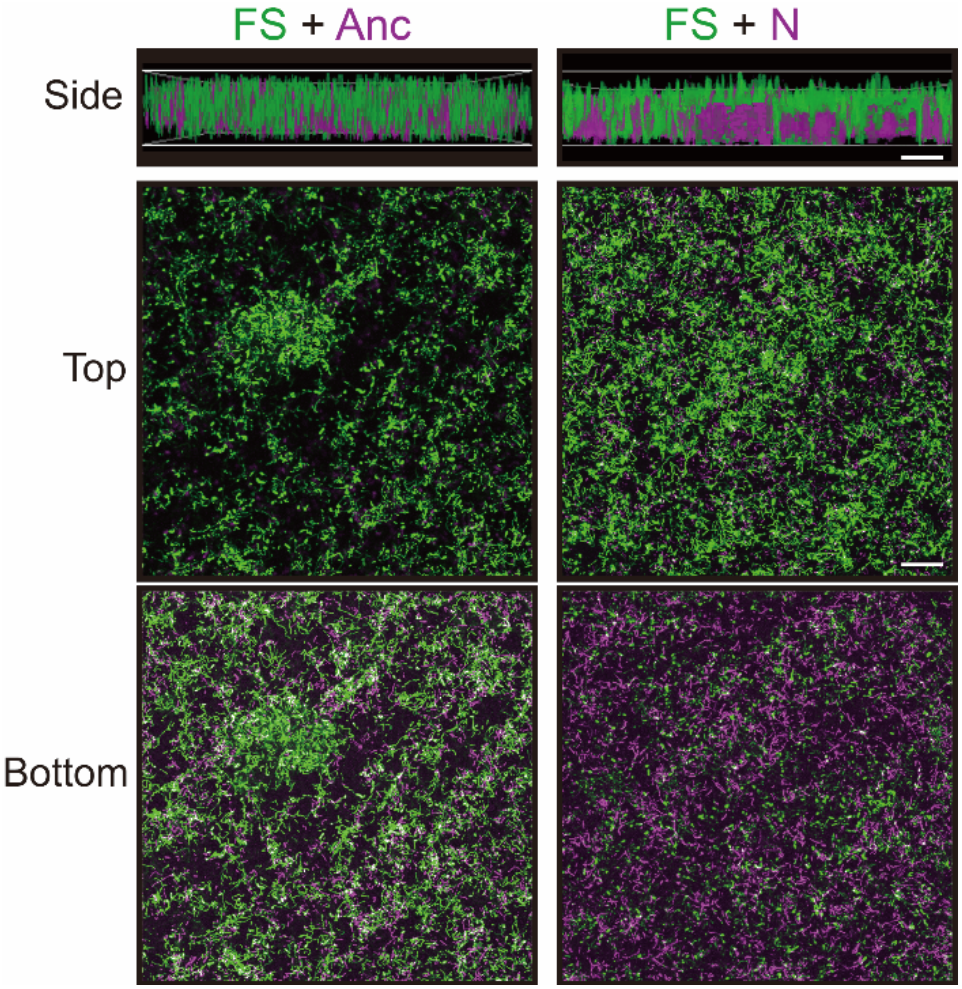
Biofilm architectures of the competitive experiments between the FS morphotype and the other two variants. Images represent the aerial view of mixed biofilms of the GFP-labelled FS morphotype and the mKate-labelled N variant or ancestor. Scale bars indicate 50 μm. The image is representative of three biological replicates.

The structural characteristics of the mixed biofilms might confer the top layer-residing FS variants significant benefits in terms of dispersal and reattachment. While only few studies have experimentally examined biofilm dispersal of Gram-positive *Bacilli*^42^, an adapted methodology was applied to monitor the dispersal of three strains into the new environment^43–45^. Briefly, mature biofilms were incubated with fresh medium, then dispersal of bacteria were quantitatively monitored over time. As expected, significantly more dispersed FS cells were present in the medium compared with the dispersal of the other two strains (Fig. S6).

### Genomic characterization of evolved isolates

To dissect the genetic alterations of the evolved morphotypes, genomic DNA of selected FS and N morphotypes from each population were extracted and the genome sequences were determined by the combination of Illumina and Nanopore sequencing (Table S1). The analysis revealed 3 to 13 SNPs compared to the previously re-sequenced ancestor genome^30^. While some of these SNPs could be found in majority of isolates (e.g. 11 SNPs were identified at least 6 or more evolved isolates), no SNP was found to be present in all isolates, highlighting the variance of evolutionary outcome. Nevertheless, no common SNP found identified in the FS variants compared to the N morphotypes. Therefore, we took advantage of the genome assemblies provided by the hybrid assembly based on the Illumina and Nanopore reads. This analysis revealed numerous genome rearrangements in the evolved morphotypes (Table S2).

Notably, compared to the ancestor and the N variants, all six FS isolates contained an insertion element in the BTB_RS26870 gene, annotated as cupin domain-containing protein. Subsequent detailed analysis revealed that this gene encodes a mannose-1-phosphate guanylyltransferase (GDP) enzyme, which belongs to the glycosyltransferase family A. In numerous organisms, GDP involves in GDP-mannose biosynthesis, which acts as the precursor for mannose residues in cell surface polysaccharides^46^. Intriguingly, mutation in the *Pseudomonas fluorescens* gene *pflu047* coding for glycosyltransferase led to modified surface lipopolysaccharides (LPS), and a fuzzy spreader morphotype^18^.

Alignment results showed that in FS morphotypes, the guanylyltransferase-encoding gene was interrupted by an insertion sequence (IS) element, annotated as IS4-like element IS231A family transposase in the genome of Bt407 (Fig. 5A). The disrupted gene located at a genetic operon comprised of 7 structural genes that are likely involved in GDP-mannose biosynthesis (Fig S7). Interestingly, comparing the FS isolates from the different evolved populations revealed that the orientation of IS element is reversible (Fig. 5A). IS element influenced genomic rearrangements are wildly observed in both natural isolates of *B. thuringiensis*^47–49^. Such property is considered to be a dynamic genomic state, which plays an important role in environmental adaptation and biological interactions of the species^50^. In the reference genome of Bt407, two copies of IS4-like element gene are present, revealing the possible plasticity of the genome and that IS element has contributed to its adaptability.

**Fig. 5.**
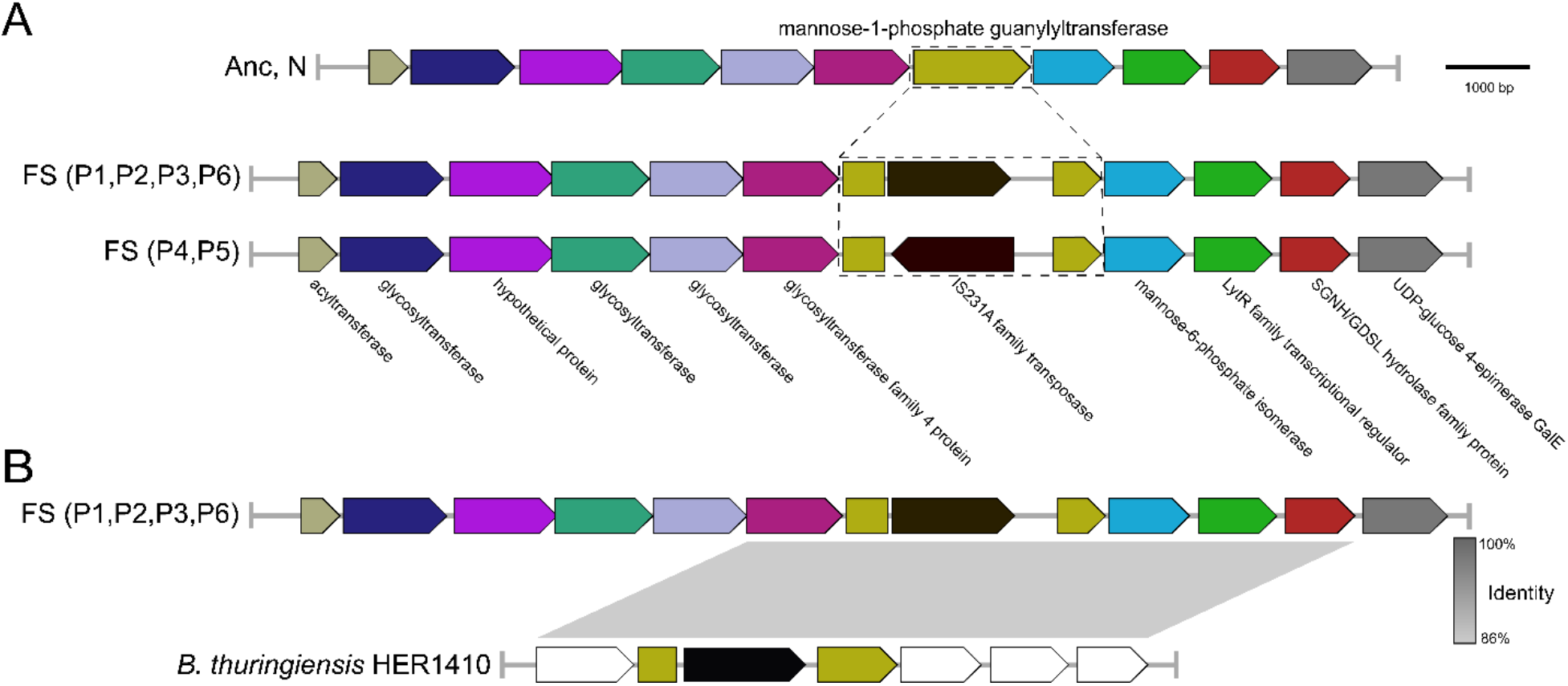
Comparative genomic characterization reveals insertion sequence plays an important role in the evolutionary adaptation of Bt407. (A) Genetic properties of the disrupted guanylyltransferase gene. (B) Identification of a similar genetic structure of the natural isolate, *B. thuringiensis* HER1410 with a guanylyltransferase disrupted by a homologous insertion sequence.

Next, to examine if such an identical insertion sequence could be detected in this gene (mannose-1-phosphate guanylyltransferase) within *Bacilli*, all complete *Bacillus* genomes available on NCBI (785 in total) were blasted against the mutated gene, the insertion sequence and the disrupted locus sequences. Overall, 20 genomes seemed to harbour the guanylyltransferase gene including one genome that contains an interrupted mannose-1-phosphate guanylyltransferase gene by a mobile element annotated as IS4 family transposase, which shares 74.6% similarity to the IS4-like element IS231A family transposase (Fig. 5B). This genome (NZ_CP050183.1) of *B. thuringiensis* strain HER1410 is known to host various bacterial phages^51^. Our *in silico* analysis, while preliminary due to the lack of direct experimental examination of strain HER1410, highlights the plasticity of mannose-1-phosphate guanylyltransferase and possible contribution to the adaption under certain environments.

### Mannose-1-phosphate guanylyltransferase influences the surface properties of Bt407

To elucidate whether the loss-of-function of *rfbM* is alone responsible for the observed FS morphotype, the complete gene was deleted in the ancestor Bt407 background by homologous recombination. The constructed mutant, Bt407Δ*rfbM*, demonstrated archetypal fuzzy colony morphotype on LB agar plates verifying that the loss-of-function mutation in *rfbM* could create FS phenotype (Fig. 6A).

**Fig. 6.**
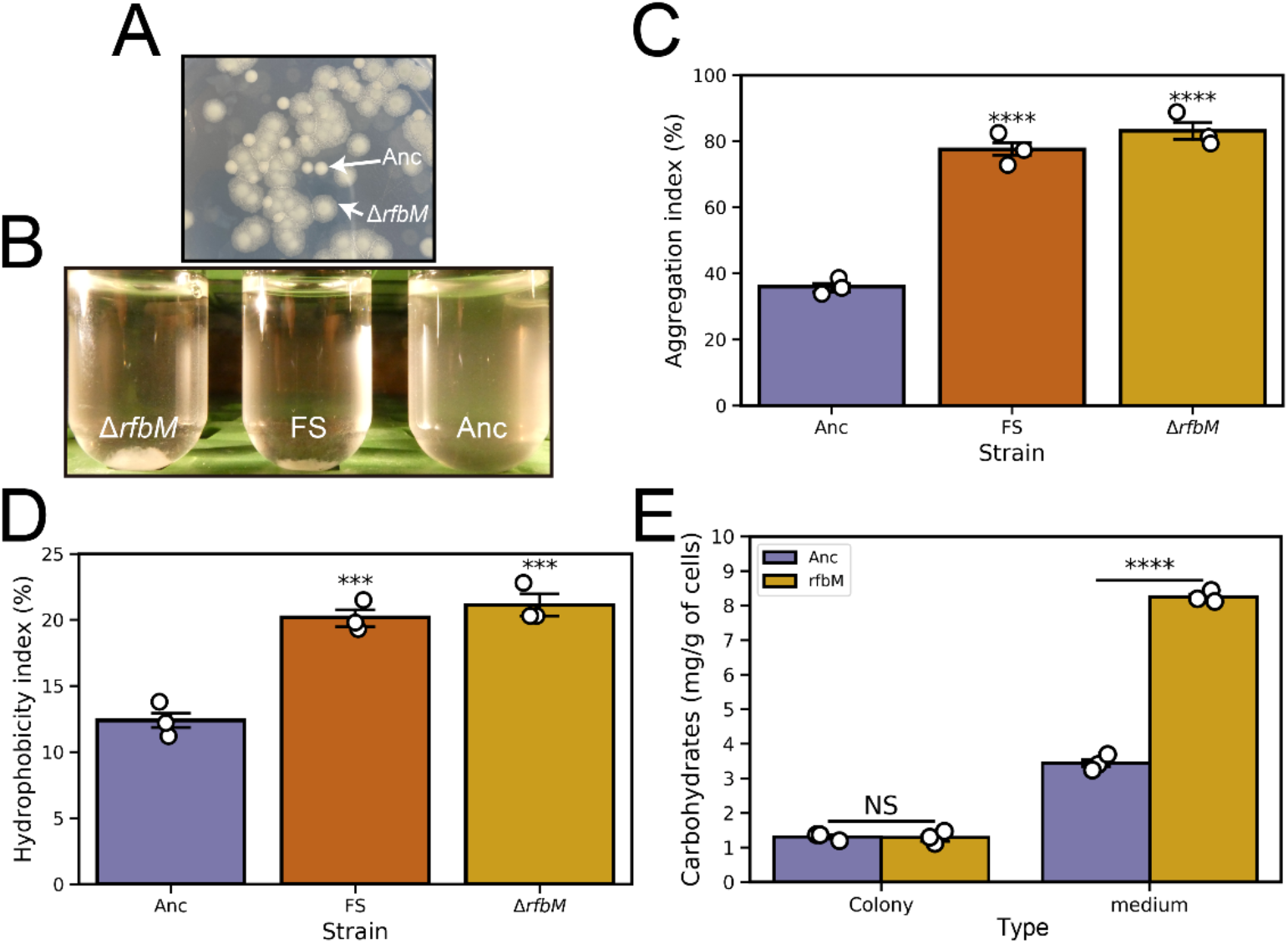
Phenotypic characterization of the mutant Bt407Δ*rfbM* strain compared with the ancestor. (A) The easily recognizable fuzzy spreader colony morphology by Bt407Δ*rfbM* suggest that disruption of the guanylyltransferase coding gene is responsible for the FS morphotype. (B) Aggregation phenotypes of Bt407Δ*rfbM* and FS variant compared with the ancestor. Aggregation (C) and hydrophobicity (D) index was characterized as described in the materials and methods (\*\*\**p* <0.001, \*\*\*\**p* <0.0001; One-way ANOVA followed by Dunnett’s multiple comparison tests between the ancestor as control and other groups). (E) Total carbohydrates determination of Bt407Δ*rfbM* and the ancestor. Asterisks indicate significant differences between each group and the ancestor (\*\*\*\**p* <0.0001; Student’s unpaired two-tailed *t* test was performed between samples within each condition). Error bars indicate standard error of the mean of independent biological samples (n = 3).

In various Gram-negative organisms, mannose-1-phosphate guanylyltransferase is involved in the biosynthesis of the capsular polysaccharide and associated with the LPS structure^52–55^. Disruption of the gene leads to the modification of the membrane elements and cell surface properties, thus affecting cell-cell interaction and biofilm architecture. According to KEGG orthology, in *Bacilli* species, mannose-1-phosphate guanylyltransferase is responsible for mannose metabolism, glycan and O-antigen nucleotide sugar biosynthesis, which also tightly regulate the surface properties. To experimentally examine whether *rfbM* contributes to physical cell-cell interactions, autoaggregation properties of FS and Bt407Δ*rfbM* were quantitatively assessed. While the parental strain demonstrated very little auto-aggregation, both the *Bt407ΔrfbM* mutant and evolved FS isolate exhibited enhanced auto-aggregation (Fig. 6B and C). In addition to the identical morphology of the colonies, this result confirmed the phenotypic consilience among fuzzy spreader and the *rfbM* deletion mutant.

Hydrophobicity is important trait that is affected strongly by cell surface structures^56–58^. Throughout this study, biofilm assays were conducted using hydrophobic objects such as nylon beads and polystyrene plates, and increased biofilm formation of FS morphotypes suggested increased hydrophobicity. MATH test was performed using the parental strain, Bt407Δ*rfbM* mutant and FS isolate to quantitatively assess the ability of bacterial adhesion to hydrocarbons. As shown in the Fig. 6D, the parental strain displayed a low level of hydrophobicity, with only 13% of cells partitioned into the hydrocarbon phase. On the contrary, higher proportion of Bt407Δ*rfbM* mutant and FS isolate partitioned into the hexadecane solvent, indicating the mutation led to a cell with increased hydrophobic cell surface.

In summary, enhanced multicellularity of FS morphotype was possibly influenced by altered characteristics of the cell surface, higher level of auto-aggregation, and hydrophobicity. More specifically, activity of the mannose-1-phosphate guanylyltransferase affect the cell surface properties and the disruption of *rfbM* by the insertion element created the observed fuzzy spreader morphotype.

## Discussion

Since the first report introducing the bead-based experiment evolution setup^24^, which primarily focused on the ecological mechanisms that sustained biofilm diversity, numerous follow-up studies have validated the robustness of this model by describing the mutational patterns^59^, genetic properties^53,60–63^, niche complementarity^64,65^, and antibiotic resistance^66,67^. Unlike other laboratory-based experimental evolution systems, this simple method is useful for modelling the complex biofilm cycle, including initial attachment, biofilm maturation, dispersal and recolonization^68,69^.

Previously, we have investigated the evolution of *B. thuringiensis* 407 biofilms on plants by repeated selection for root-associated biofilm cycles^30^. The bead biofilm model provides a simpler, abiotic selection system to reveal how adaption to the biofilm life cycle influences bacterial evolution. Although genetic and phenotypic differentiation were already observed with the plant-adapted isolates, the bead-based model created highly predictable multicellular phenotypic differentiation. Namely, a morphologically distinct variant was observed in each population that was distinct from the ancestor, exhibiting large colony and fuzzy appearance. Adaptive diversification have pivotal contribution to the evolution of bacterial community, mostly influencing biological diversity^70^. Spatially structured environments are a critical factor that facilitates such variation that allows competition of variant morphotypes and emergence of newly evolved niche-specialists, so called ecotypes. In addition to the bead model and static microcosms that were used for Gram-negative bacteria ^14,71,72^, diversification was also observed among Gram-positives, e.g. laboratory evolution of *B. subtilis* in static, shaking, and pellicle conditions^25–27^. Using a host associated setup, Blake *et al*. illustrated how *B. subtilis* diversified into several morph variants when evolving in plant root-associated biofilms^38^.

Here, a unique Fuzzy Spreader was identified that displayed specific cellular behavior phenotypes, including altered swimming, surface spreading, biofilm formation in addition to overall enhancement of bead colonization. Interestingly, FS variants formed dendritic or branched patterns on EPS or Trb media with 0.7% agar. In *B. cereus*, dendrite colony formation was induced by low-nutrient conditions like EPS and was connected to the production of bio-surfactant compounds, which was negatively regulated by PlcR regulon in *B. cereus* ATCC 14579^73^. The extensive dendritic pattern of FS variants suggests an efficient translocation strategy, which may contribute to colonizing and surviving in a new environment. While the FS variant clearly exhibited distinct differentiation properties compared to the ancestor and the evolved N variant, both FS and N morphotypes from P1 demonstrated elevated Congo red binding property, suggesting higher production of biofilm matrix. Enhanced biofilm matrix production in addition to potentially increased bio-surfactin production might synergistically promote colony expansion of the FS variants on 0.7% agar medium. Indeed, sliding in *B. subtilis* depends on the production of both surfactant compounds as well as exopolysaccharides^74–78^. Surfactin facilitates colony expansion by reducing the friction between cells and their substrate. In *B. subtilis*, cell collectives organized as bundles, termed as van Gogh bundles, which depends on the synergistic cooperation of surfactin- and matrix-producing cells^78^. Hence, whether FS variants display enhanced sliding or flagellum dependent swarming requires further studies using directed microscopy and mutant generation.

Intriguingly, FS variants exhibited distinct biofilm development in LB and EPS medium. The dissimilar biofilm on these two media might be caused by different amount and composition of nutrients, aggregation of FS variants is for example induced by nutrient-limited environments. Multicellularity confers bacteria distinct characteristics that can be advantageous or disadvantageous depending on the niche they occupy; therefore, it is interesting to determine what evolutionary factors promote multicellular groups instead of dispersed unicellular populations? Series of evolution experiments have been developed both *in vivo* and *in silico* to demonstrate how multicellularity can evolve^23,72,79–83^. For instance, multicellularity evolved 2.5 billion years ago in cyanobacteria^84^. Cyanobacteria were shaped by various environments and diversified through distinct evolutionary paths, resulting in both unicellular and multicellular forms that are found also today^85,86^. This implies that multicellularity readily evolves through adaptive diversification, microbes could diversify into certain pheno- and genotypic variants in specific niches.

One of the emerging functions of multicellularity is the improved performance of the group compared to the undifferentiated population. Cooperative interaction could be indeed detected between the FS and N morphotypes when forming mixed biofilm compared with the biofilms created by one of evolved type or the ancestor strain alone. Further, cooperative and competitive cell-cell interactions shape spatial assortment of genotypes within a biofilm community^40^. Our CLSM analysis of mixed biofilms revealed that FS variants is distributed more evenly when cocultivated with evolved N variant compared with the aggregated distribution when combined with the ancestor, potentially suggesting the emergence of a synergistic interaction between the evolved morphotypes. Moreover, the high competitiveness of FS variants in the presence of ancestor suggests increased translocation ability of FS variants, which might correlate with the increased surface spreading. Individual-based modeling has previously suggested that ability of extracellular polymer production confers a strong competitive advantage within mixed genotype biofilms^87^. Interestingly, morphologically similar variants similar to the FS type observed in this study was also identified by Rainey et al.^14^ after evolution of *P. fluorescens* in a static microcosm allowing spatially heterogeneous diversification. It has been suggested that fuzzy-spreader morphotype of *P. fluorescens* exhibits niche specialization by forming submerged aggregates at the bottom of tube^88–90^. Like wrinkly-spreader, fuzzy-spreader displays a multicellular phenotype including increased biofilm formation ability^72^. However, the selective advantage of the fuzzy-spreader morph was not clearly clarified while wrinkly-spreader was comprehensibly examined by Rainey group^35,91–93^. These studies laid a foundation for subsequent pheno- and genotypic characterization of fuzzy-spreader^18^, demonstrating that fuzzy-spreader follows an ecological cycle of forming cellular rafts, collapsing into the bottom of the vial and reforming the rafts in the microcosms, which displays analogy to the repetitive life cycle of FS morphotype in this study. The genetic dissection of fuzzy-spreader of *P. fluorescens* identified a gene, *fuzY*, which encodes a β-glycosyltransferase that is predicted to modify O antigens of lipopolysaccharide (LPS)^18^. Mutation in *fuzY* leads to cell flocculation. Essentially, LPS defects were associated with stronger adhesion of cells both to abiotic surfaces and among each other. Similarly, the FS morphotypes of *B. thuringiensis* harbored insertion elements within a gene encoding a mannose-1-phosphate guanylyltransferase. The disrupted *rfbM* gene is located in an operon that is predicted to be responsible for polysaccharides biosynthesis. In *Burkholderia* and *Pseudomonas* genera, disruption of LPS biosynthesis may improve biofilm properties such as adhesiveness, cohesiveness, and viscoelasticity^94,95^. Even though LPS has crucial function in Gram-negatives, cells of Grampositive bacteria are not encased by a lipopolysaccharide layer, rather having a complex layered structure including peptidoglycan, peptides and amino acids^96,97^. Nevertheless, in Gram-positive bacteria, mannose-1-phosphate guanylyltransferase is involved in the biosynthesis of capsular polysaccharide (CPS), which are polysaccharides associated with the cell surface^98^.

Cellular hydrophobicity and auto-aggregation are highly dependent upon non-polar functional elements, surface protein, pilus and capsules. It has been previously emphasized that CPS plays an essential role in determining biofilm size by various mechanisms such as inhibiting continual growth of matured biofilms via quorum sensing^57,99,100^. Variation of such membrane-associated structure has been shown to directly influence bacterial surface properties, thus cell-cell and cellsurface interactions that influence biofilm architecture in numerous bacteria^101,102^. In *Porphyromonas gingivalis*, mutation in a glycosyltransferase gene was responsible for enhanced biofilm formation and hydrophobicity because of the adjustment of cell surface characteristics^103^.

### Conclusions

Our work provides the first comprehensive assessment of evolutionary diversification of *B. cereus* associated to abiotic biofilms. The findings reported here shed new light on previously unknown adaptive strategy of *B. cereus* by diversifying into phenotypically distinct morphotypes in response to biofilm life cycles. FS variant exhibited distinct multicellular phenotypes with specifically increased biofilm development. FS variant retained the highest fitness and acted as a generalist that balances biofilm formation and dispersal due to a disrupted guanylyltransferase-coding gene. Genomic characterization of FS morphotypes revealed parallel genomic plasticity via displacement of an insertion element that promoted evolutionary adaptation. Multicellularity of generalists during selection within a biofilm lifecycle could be a relevant evolutionary adaptation in environmental niches such as plants and soil, in which the biofilm lifecycle is suggested to be required for survival of bacteria.

## Materials and Methods

### Bacterial strains and growth conditions

The ancestral (wide-type) strain used in this study is *B. thuringiensis* 407 (Bt407 Cry-, commonly referred as Bt407). Table 1 includes all bacterial strains, plasmids and primers used in this study. *Escherichia coli* XL1-Blue was used for molecular cloning experiments. Routinely, bacterial strains including *B. thuringiensis* and *E. coli* were cultured in lysogeny broth (LB-Lennox, Carl Roth; 10 g/L tryptone, 5 g/L yeast extract and 5 g/L NaCl) plates solidified with 1.5% agar or stored at −80 °C with 28% glycerol added to an overnight culture. When required, concentrations of antibiotics were used as indicated: kanamycin (50 μg/mL), ampicillin (100 μg/mL), erythromycin (5 μg/mL), and tetracycline (10 μg/mL). X-gal (5-bromo-4-chloro-3-indolyl-β-D-galactopyranoside) was used at 40 μg/mL. For all relevant assays, the incubated cultures were vortexed vigorously to ensure proper disruption of any cellular aggregates.

**Table 1.**
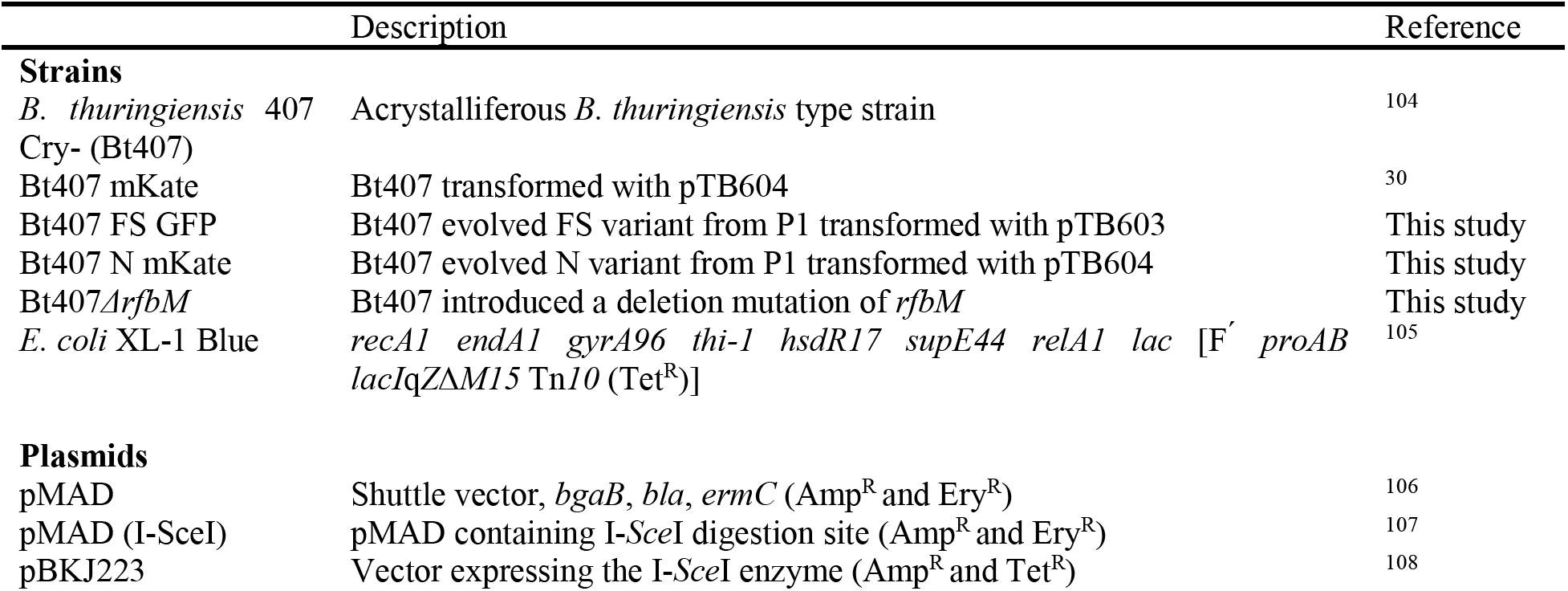
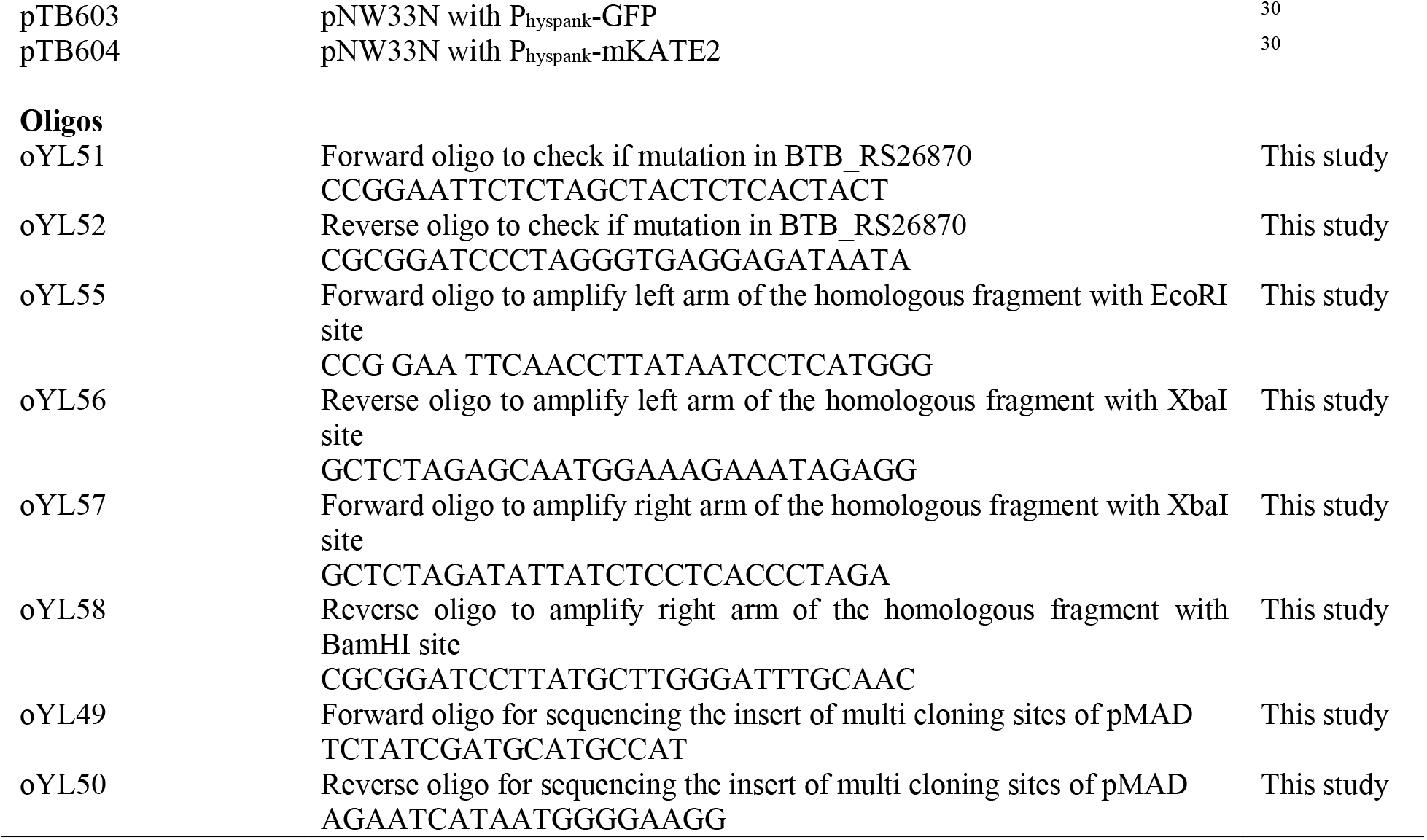
Information of strains, plasmids and oligos used in this study

### Experimental evolution

Experimental evolution setup was applied as previously described with modifications^24^. A homogenous colony of Bt407 ancestor was inoculated into a 24-well microtiter plates containing a nylon bead floating in EPS medium^109^, which was low-nutrient medium specifically for *B. cereus* biofilm formation (7 g/L K_2_HPO_4_, 3 g/L KH_2_PO_4_, 0.1 g/L MgSO_4_·7H_2_O, 0.1 g/L (NH_4_)_2_SO_4_, 0.01 g/L CaCl_2_, 0.001 g/L FeSO_4_, 0.1 g/L NaCl, 1 g/L glucose, and 125 mg/L yeast extract). The plates were sealed with parafilm and incubated at 30°C and shaking at 90 rpm. After 24 h, the colonized bead was transferred into a fresh EPS medium containing two uncolonized, sterile beads. Around every three transfers, biofilm was dispersed from one of the newly colonized beads using rigorous vortexing and subjected to standard biofilm productivity analysis that is based on determining the total cell number. The whole experiment included six individually evolved populations and additional six planktonic bacterial cultures as control without beads. The regression analysis was performed towards all replicates with the timeframe from the beginning to transfer 31 excluding an outlier (transfer 25).

### Genetic manipulations

Electroporation of *B. thuringiensis* and DNA extraction were performed as standard procedures^104,110^. Restriction enzymes, T4 DNA ligase, and Phusion High-Fidelity DNA Polymerase were purchased from Thermo Scientific. DNA fragments were purified by using NucleoSpin Gel and PCR Clean-up kits (Macherey-Nagel). Oligos were synthesized by TAG Copenhagen A/S and DNA sequences were sequenced at Eurofins Genomics. For *E. coli*, standard molecular was applied according to standard protocols^111^.

The deletion mutant of *B. thuringiensis* was constructed by homologous combinations using a mark-less replacement method introduced by Janes and colleagues^108^. Briefly, the flanking homologous fragments of approximately 700bp were amplified and cloned into the shuttle vector pMAD^106^ with an additional I-SceI restriction site^107^. The plasmid carrying the flanking homologous fragments was electroporated into Bt407 to obtain blue transformants on LB agar plates containing erythromycin and X-gal. Facilitation of the integration into the chromosome was conducted as previously described^108^. Subsequently, pBKJ223, which encodes I-SceI restriction enzyme, was transformed to induce double-stranded break at the chromosome thus promoting a second recombination event^112^. Finally, White colonies represented as those that have lost erythromycin resistance were selected and genomic DNA was extracted. Mutation was verified by PCR and Sanger sequencing.

### Fitness assays of bead biofilms

Fitness was determined for both relative and competitive biofilm assays, where the ratio of Malthusian parameters (*m*) was calculated described by Lensiki *et al*.^37^. Bacteria were recovered and plated onto agar plates and calculated as *m* = ln(*N*_1_/*N*_0_) where *N*_1_ and *N*_0_ represented colony forming units counted in the end and at the start of the assays, respectively.

### Phenotypic characterization of genetic variants

#### Congo red indicator assay

10 μl overnight grown bacterial culture was spotted onto LB agar plate (1.5%) supplemented with 40 μg/ml of Congo red (CR) and 20 μg /ml of Coomassie brilliant blue dyes. Bacterial colonies were grown at 30°C, after which images were taken using Panasonic DC-TZ90 camera.

#### Surface motility assay

10 μl overnight culture (approx. 10^6^ CFU) was spotted onto the center of 0.7% agar plate containing a nutrient medium and incubated at 30°C for up to 72 h. Colony images were recorded with a Panasonic DC-TZ90 camera. EPS agar plates were made with EPS medium supplemented with agar as described^73^. TrB (1% tryptone-0.5% NaCl) medium was used as described previously^113^.

#### Biofilm visualization

Bacterial strains were inoculated in rich LB medium LB and incubated for 24h at 30°C. Biofilms forming on the tube walls were imaged with a Panasonic DC-TZ90 camera. Biofilm dispersal and CV staining: Dispersal was quantified according to previous report^45^. Bacterial cultures were incubated in EPS medium for 24 h to form mature biofilms in 24-well microtiter plates, followed by gentle wash of unattached cells. One-mL fresh EPS medium were added into the plates, which was then agitated vigorously for 1 h and 10h represented as weak and strong disturbance, respectively. Remaining biofilms were stained by crystal violet (1%) and solubilized by ethanol, after which A490 was documented.

### Microscopy

For bright-field images of bacterial pellicles and colonies, Axio Zoom V16 stereomicroscope (Carl Zeiss) was used, which equipped with a Zeiss CL 9000 LED light source, a PlanApo Z 0.5 × objective, and AxioCam 591 MRm monochrome camera (Carl Zeiss, Jena).

Confocal laser scanning microscopy (CLSM) was conducted as described previously^114^. For surface attached biofilms imaging, bacterial cultures were incubated in high content imaging plate (Corning, New York) to form mature biofilms as described above. Afterwards biofilms were washed with sterilized ddH2O twice to remove non-attached aggregates. CLSM images were obtained using a 63 × /1.4 OIL objective. Fluorescent excitation was conducted with the argon laser at 488 nm and the emitted fluorescence was acquired at 484-536 nm and 567-654 nm for GFP and mKate, respectively. Z stack series of biofilms were obtained with 1 μm steps and stacked images were merged using ImageJ software.

### Whole genome sequencing and hybrid assembly

A 4-mL aliquot of bacterial cultures of 12 evolved isolates (representative FS and N variants from 6 evolved populations) plus the ancestral strain was centrifuged and genomic DNA was extracted using GeneMATRIX Bacterial Genomic DNA Purification Kit (EURx Ltd, Poland). Paired-end libraries were prepared using the NEBNext^®^ Ultra™ II DNA Library Prep Kit for Illumina sequencing. Illumina NextSeq sequencer was used to generate paired-end fragment reads using TG NextSeq^®^ 500/550 High Output Kit v2 (300 cycles).

Long-read libraries were constructed using Rapid Barcoding Sequencing kit (SQK-RBK004) and long-reads were generated on MinION Mk1B (Oxford Nanopore Technologies). Raw sequences were base called using MinKNOW sequencing software (Oxford Nanopore Technologies). Preprocessing was performed using AdapterRemoval and Filtlong for short and long reads, respectively.

Subsequently, reads of each isolate were hybrid assembled with short reads from Illumina sequences by using Unicycler^115^. Assembled genomes of evolved isolates and the ancestor was aligned using MAFFT^116^. Finally, assemblies were visualized in CLC workbench and annotated by Prokka. Easyfig was used for visualization of the genomic alignment^117^.

The assembled genomes were compared to *B. thuringiensis* 407 genome (GenBank accession no. CP003889.1), after extraction of previously mapped SNPs found in the ancestor used in this study^30^. SNPs were identified between the reference genome (GenBank accession no. CP003889.1) and the assembled contigs using Snippy (GitHub - tseemann/snippy at v4.6.0). Identified mutations are included in TableS1. Raw sequencing data has been deposited to the NCBI Sequence Read Archive (SRA) database under BioProject accession number: PRJNA757963.

Profiling of *Bacillus* genomes for disrupted *rfbM* genes was done by downloading all completed *Bacillus* genomes from NCBI using ncbi-genome-download (https://github.com/kblin/ncbi-genome-download) and searching for the intact gene, the IS and the interrupted gene using blastn.

### Cell surface characterization

Adhesion to hydrocarbons (BATH) assay was conducted as previously^56^. Briefly, cells were washed and resuspended in PBS buffer to remove culturing medium. Bacterial suspension was then adjusted to optical density (600nm) of 0.25 ± 0.05 (recorded as *A*_0_), representing the standard number of bacteria (10^7^–10^8^ CFU/ ml). Then, an equal volume of Hexadecane (Sigma-Aldrich) was added. The two-phase liquid system was mixed completely by vortexing vigorously for 10 min, following by a 1 h of incubation at room temperature allowing stratification. The optical density (600nm) of the aqueous phase was measured as *A*. The hydrophobicity, represented as adhesion to hydrocarbons, was calculated across three different replicates according to the formula: 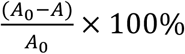.

For auto-aggregation analysis, liquid cultures were monitored as previously reported^118^. Briefly, glass tubes containing 5 mL of EPS were inoculated with 24-hour-old colonies and cultured overnight in a 37°C incubator and shaking at 220 rpm. Bacterial cultures were then settled for 10 h at 4°C. Auto-aggregation was quantified by measuring the cell density changes before and after settlement as following: 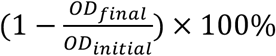. Auto-aggregation index was calculated from triplicates.

### Carbohydrates quantification

Collection of exopolysaccharide was applied according to previous work^**119**^. Briefly, strains were cultivated at 30°C for 48h on EPS agar (1.5%) plates, or in EPS liquid medium with shaking at 220 rpm. Bacterial colonies were suspended in one mL of 0.9% NaCl buffer and subjected to vigorous sonication (5 × 12 pulses of 1 s with 50% amplitude; Ultrasonic Processor VCX-130, Vibra-Cell, Sonics, Newtown). Liquid bacterial cultures were treated with the same sonication process. Bacterial biomass was separated by centrifugation (10 min at 12000g) and supernatant was collected. Exopolysaccharide content of samples was quantified by using the phenol-sulfuric acid method described previously^120^. Standard curve was constructed using diluted glucose solution (*y* = 19.773*x* + 0.0827, R^2^ = 0.9984).

### Statistical analysis

Unless indicated otherwise, all experiments were performed with at least three biological replicates. Statistical analysis of bacterial traits comparison between evolved isolates and the ancestor was analyzed and illustrated using Python 3.8 with Statsmodels packages or Graphpad 8. For test statistically significant differences between the means of three or more independent groups, one-way ANOVA analysis was carried out, followed by Dunnett’s post-hoc analysis. For comparing the means of two groups, student’s unpaired two-tailed *t* test was performed.

## Acknowledgments

Y.L. was supported by a Chinese Scholarship Council fellowship. G.M. was supported by the 685 Lendület-Programme of the Hungarian Academy of Sciences (LP2020-5/2020). Funding from Novo Nordisk Foundation for the infrastructure “Imaging microbial language in biocontrol (IMLiB)” (NNFOC0055625) is acknowledged. The position of M.L.S. is financed by the Danish National Research Foundation (DNRF137) for the Center for Microbial Secondary Metabolites.

