## Supplementary material for "Adaptation and phenotypic diversification of *Bacillus thuringiensis* 407 biofilm are accompanied by a fuzzy spreader morphotype": Fig S1 to S7

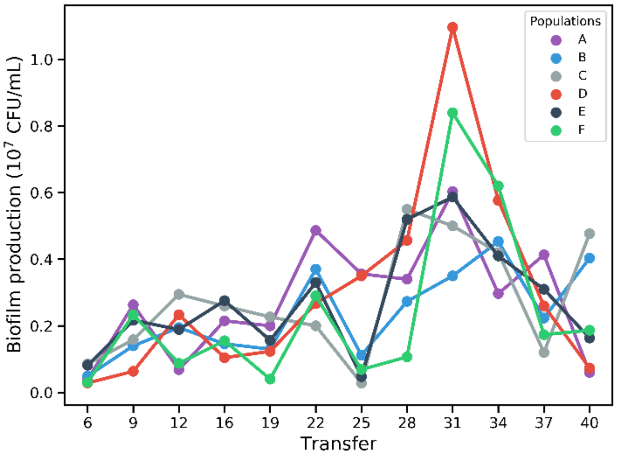


**Fig. S1** Biofilm productivity of bead-colonized Bt407 (Cry$^-$) of all six evolved populations are shown at roughly every 3$^{th}$ transfer.

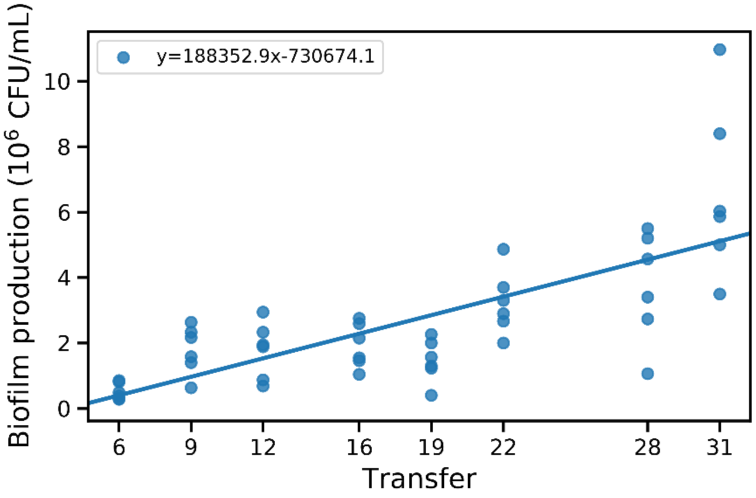


**Fig. S2** Fit linear regression of the average biofilm productivity of bead-colonized Bt407 ($Cry^-$) of all six evolved populations.

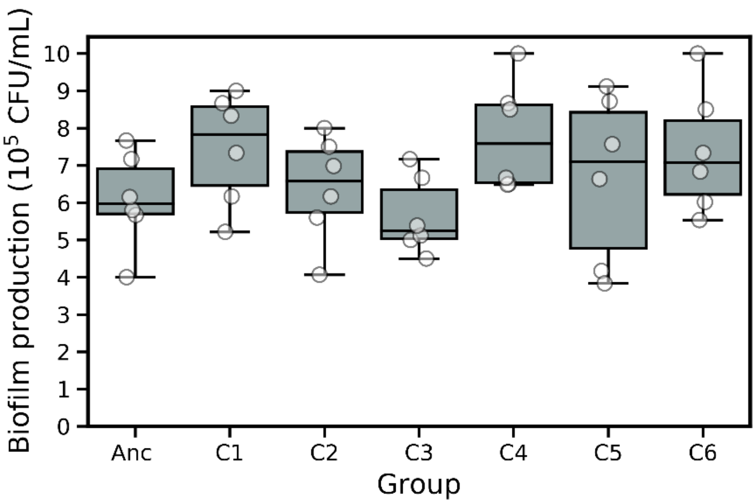


**Fig. S3** Biofilm productions of six control populations compared with the ancestor. Boxes indicate Q1–Q3, lines indicate the median, and bars span from max to min. Error bars indicate the standard error of six biologically independent replicates.

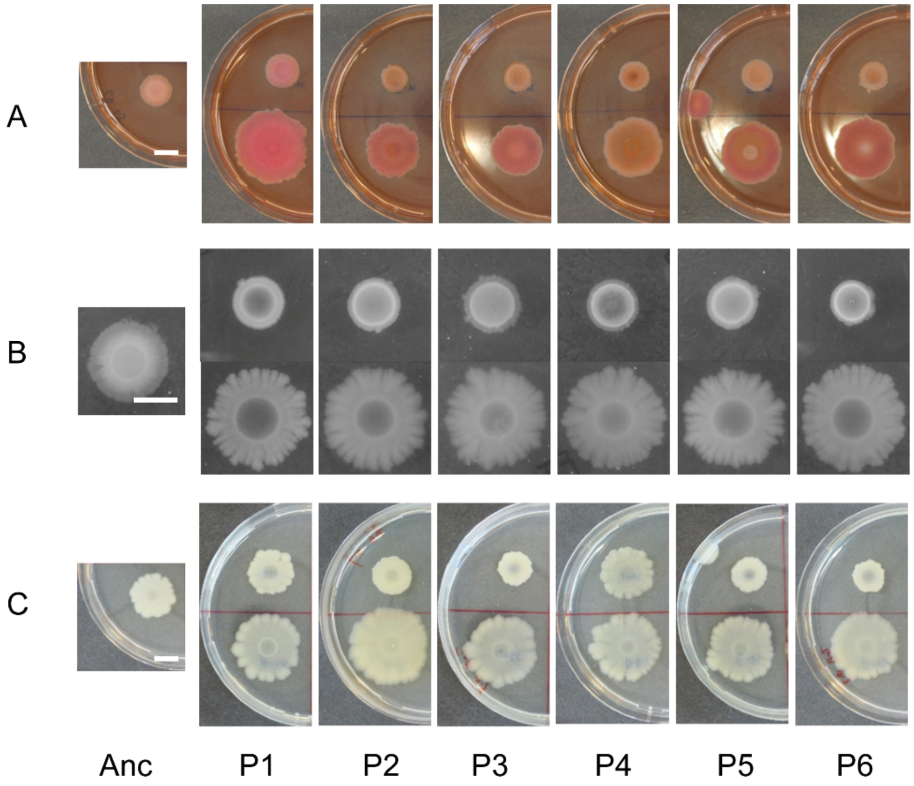


**Fig. S4** Screened phenotypical characterization of evolved morph variants from all six evolved populations compared with the ancestor, including Congo red uptake colony morphologies (A), swarming motility on EPS (B) and TrB (C) medium contain 0.7% agar. Top and bottom images represent FS and N evolved variants, respectively. Scale bars indicate 10 mm for all panels.

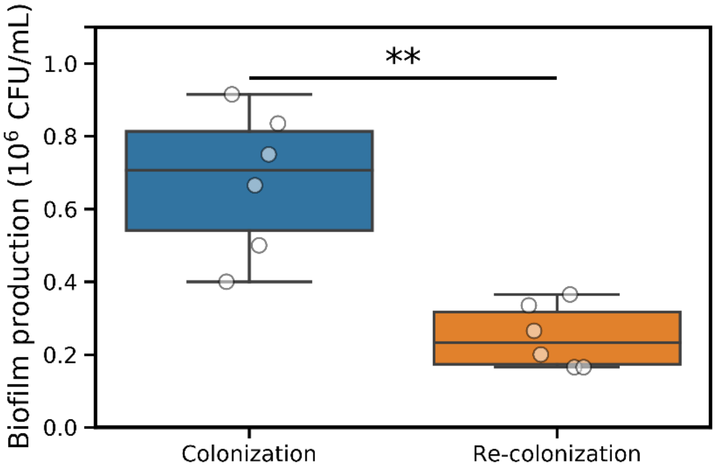


**Fig. S5** Statistical different biofilm productions of colonization and re-colonization in the ancestor strain. Boxes indicate Q1–Q3, lines indicate the median, and bars span from max to min. Error bars indicate the standard error. Asterisks indicate significant differences between each group and the ancestor (***p* <0.01; two tailed t-test with Welch's corrections)

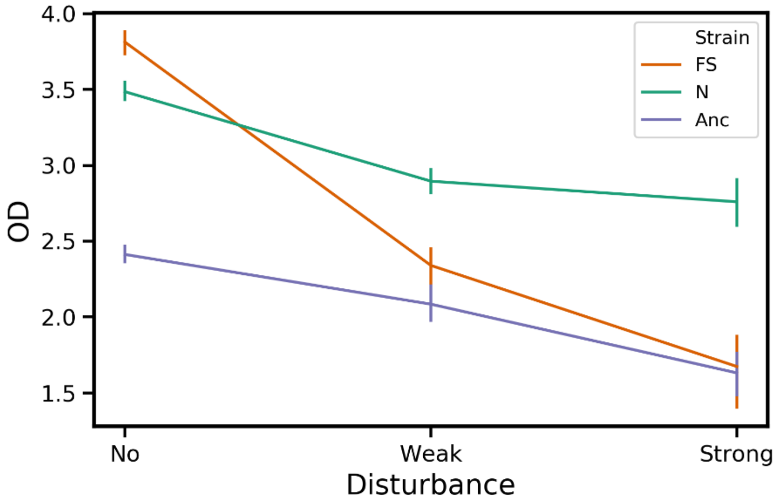


**Fig. S6** Bacterial dispersal ability was quantified with FS, Normal morph variants and the ancestor (n=10). Bacterial cultures were incubated in 24-well microtiter plates. OD as biofilm formation in the wells by crystal violet staining was monitored for three different groups. Plates were vigorously shaking on an orbital shaker (220 rpm) for 1 h and 10 h represented as weak and strong disturbance to induce biofilm dispersal, followed by staining of the biofilms using crystal violet.

A

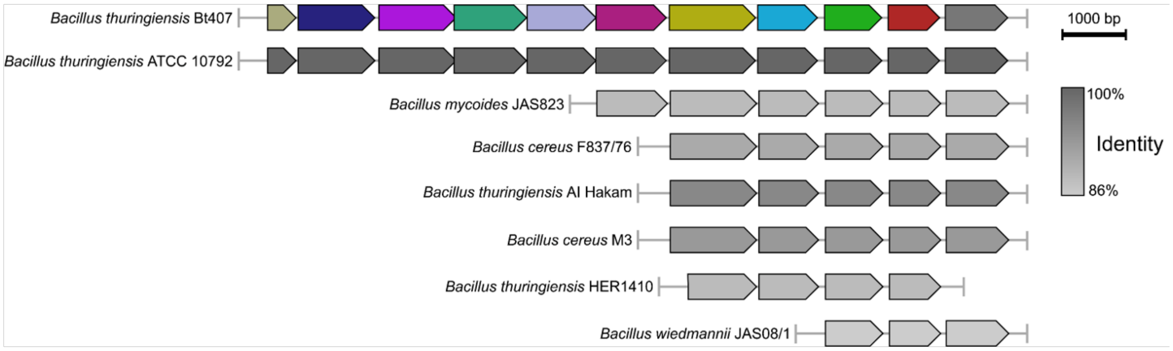


B

| Operon | IdGene | Type | COGgene | PosLeft | postRight | Strand | Function |
|---|---|---|---|---|---|---|---|
| 1 | | | | | | | |
| | MNDAOHKI_00011 | CDS | COG1316 | 12656 | 13570 | - | [K] Transcriptional regulator |
| | MNDAOHKI_00012 | CDS | COG1482 | 13682 | 14656 | - | [G] Phosphomannose isomerase |
| | MNDAOHKI_00013 | CDS | COG0836 | 14666 | 16039 | - | [M] Mannose-1-phosphate guanylyltransferase |
| | MNDAOHKI_00014 | CDS | COG0438 | 16087 | 17211 | - | [M] Glycosyltransferase |
| | MNDAOHKI_00015 | CDS | COG0438 | 17208 | 18302 | - | [M] Glycosyltransferase |
| | MNDAOHKI_00016 | CDS | COG0438 | 18313 | 19464 | - | [M] Glycosyltransferase |
| | MNDAOHKI_00017 | CDS | COG3872 | 19436 | 20665 | - | [R] Predicted metal-dependent enzyme |
| 2 | | | | | | | |
| | MNDAOHKI_00018 | CDS | COG0438 | 20730 | 21947 | - | [M] Glycosyltransferase |
| | MNDAOHKI_00019 | CDS | COG0110 | 21984 | 22439 | - | [R] Acetyltransferase (isoleucine patch superfamily) |
| | MNDAOHKI_00020 | CDS | COG2244 | 22761 | 24044 | - | [R] Membrane protein involved in the export |
| | MNDAOHKI_00021 | CDS | COG0677 | 24255 | 25511 | - | [M] UDP-N-acetyl-D-mannosaminuronate dehydrogenase |
| | MNDAOHKI_00022 | CDS | COG2148 | 25531 | 26211 | - | [M] Sugar transferases involved in lipopolysaccharide synthesis |
| | MNDAOHKI_00023 | CDS | COG1210 | 26230 | 27111 | - | [M] UDP-glucose pyrophosphorylase |

**Fig. S7** (A) Comparative genomic characterization of disrupted genetic region of Bt407 compared with the homologous regions of other species in *B. cereus* group. (B) Prediction operon was conducted surrounded the disrupted gene as red rectangle indicates.
